# Aurora A kinase activation contributes to the fibrotic phenotype in Systemic Sclerosis through primary cilia shortening

**DOI:** 10.64898/2026.03.13.711548

**Authors:** Rebecca Wells, Begoña Caballero Ruiz, Panji Mulipa, Alex J. Timmis, Maria E. Teves, John Varga, Francesco Del Galdo, Rebecca L. Ross, Natalia A. Riobo-Del Galdo

## Abstract

**Background:** Systemic sclerosis (SSc) is a severe autoimmune disease characterised by progressive fibrosis driven by fibroblast activation. Primary cilia, key hubs for profibrotic signalling, are markedly shortened in SSc fibroblasts, but the mechanisms underlying this phenotype remain unclear. This study aimed to define the signalling pathways responsible for primary cilia shortening and fibroblast activation in SSc.

**Methods:** Primary dermal fibroblasts from SSc patients and healthy controls were analysed for cilia incidence and length by immunofluorescence, profibrotic marker expression by qPCR, and contractility using gel contraction assays. Cells were treated with TGFβ1 and pharmacological inhibitors targeting AURKA, HDAC6, ROCK2, and Smad3 signalling. CAV1-silenced fibroblasts were used as an in vitro model of SSc.

**Results:** Maintenance of the constitutively short primary cilia phenotype in SSc fibroblasts did not require active TGFβ signalling. However, TGFβ1 induced reversible cilia shortening in healthy fibroblasts and further shortened cilia in SSc fibroblasts to a similar final length, mediated by Rho/ROCK2 rather than canonical Smad3-dependent signalling. Constitutive cilia shortening in SSc was driven by aberrant AURKA activity upstream of HDAC6, promoting ciliary disassembly. Pharmacological inhibition of AURKA or HDAC6 selectively elongated cilia in SSc fibroblasts, reduced profibrotic marker expression, and abrogated fibroblast contractility. CAV1-silenced fibroblasts similarly exhibited constitutive cilia shortening that was reversed by AURKA inhibition without affecting healthy cells.

**Conclusions:** Aberrant activation of the AURKA/HDAC6 axis maintains short primary cilia and promotes fibroblast activation in SSc. These findings reveal a mechanistic link between cilia morphology and fibrosis and identify AURKA as a potential therapeutic target for SSc-associated tissue remodelling.

## INTRODUCTION

Systemic Sclerosis (SSc) is a chronic autoimmune connective tissue disease characterized by widespread fibrosis, vascular abnormalities, and immune system dysregulation [1]. The hallmark of SSc is excessive collagen deposition, which leads to thickening and hardening of the skin and organ dysfunction. SSc has the highest mortality amongst rheumatic autoimmune diseases, especially in patients developing interstitial lung disease and pulmonary artery hypertension [2]. Fibrosis is the result of excessive production of extracellular matrix by myofibroblasts. These highly contractile matrix secreting cells, arise by transdifferentiating of resident connective tissue fibroblasts or other cell types including endothelial cells, keratinocytes, adipocytes, fibrocytes and pericytes in response to multiple stimuli [3]. The prototypical driver of myofibroblast activation is Transforming Growth Factor β (TGFβ); however, Hedgehog (Hh), Platelet-derived Growth Factor (PDGF) and Wnt signalling have also been shown to promote the myofibroblast phenotype in SSc [4].

Interestingly, the cellular receptors and key mediators of those profibrotic signals have been shown to localise to the primary cilium [5] [6] [7, 8]. However, to date little is known regarding the cell type-specific regulation and role of primary cilia in fibrosis. The primary cilium is a solitary, non-motile, microtubule-based organelle that protrudes from the surface of most vertebrate cells during interphase and serves as a cellular antenna for chemical and mechanical signals [9]. The primary cilium axoneme consists of nine microtubule doublets arranged in a ring without a central pair that extend from the basal body, which is derived from the mother centriole of the centrosome. The length of the primary cilium is under tight regulation with a dynamic balance between cilia elongation (assembly) and shortening (disassembly) [10]. The organelle is assembled during cell cycle arrest (G0/G1 phase) and is resorbed prior to mitosis, reflecting its dynamic regulation in coordination with cell division [9]. Maximal cilia length is achieved during G1, followed by two cycles of cilia resorption at the G1/S and the G2/M transitions prior to centriole duplication and formation of the mitotic spindle [10]. The first wave of cilia resorption is regulated by the Aurora A kinase (AURKA)/ Neural precursor cell expressed developmentally down-regulated protein 9 (NEDD9)-Histone Deacetylase 6 (HDAC6) cascade, while the second wave is mediated by the Pericentriolar Material 1 (PCM1)-Polo-like Kinase 1 (Plk1)-(Kif2A/Dzip1/HDAC6) cascades [10]. During S and G2 phases, the Nek2-Kif24 cascade inhibits re-ciliation [11]. Following mitosis, the process of ciliogenesis generates primary cilia of original length, which varies between 1 and 10 μm depending on the cell type, and reflects the balance between assembly and disassembly dynamics [12]. Ciliary disassembly can also be mediated by other mechanisms, such as cilia remodelling and cilia deconstruction [13].

In view of the predicted role of primary cilia in regulating cellular signalling elicited by fibrogenic factors, we have recently endeavoured to characterize the expression and function of primary cilia in SSc and fibrosis [14, 15]. We noted that under in vitro culture conditions, fibroblasts isolated from patients with different fibrotic conditions, including SSc, are characterised by shorter primary cilia compared to anatomic site-matched fibroblasts isolated from healthy subjects [16]. Moreover, we noted that forced elongation of primary cilia was accompanied by reduced expression of pro-fibrotic markers, suggesting that the short cilia phenotype might have a pathogenic role in fibrosis. Those findings sparked further interest in understanding the mechanisms that underlie the ciliary phenotype of activated fibroblasts.

In this study, we further characterised the molecular mechanisms leading to short cilia phenotype of SSc dermal fibroblasts. Our findings indicate that the short cilia phenotype of SSc cells is stable, and dependent on AURKA cilia-disassembly path. Importantly, inhibition of AURKA activation rescues the short cilia phenotype in SSc fibroblasts and reduces expression of fibrotic markers and cell contractility. Therefore, our results highlight the pathogenic link between primary cilia length and fibroblast activation in SSc and suggest a therapeutic opportunity by repurposing AURKA inhibitors.

## RESULTS

### Short cilia phenotype of SSc patient fibroblasts is not maintained by canonical TGFβ signalling

We have recently reported that fibroblasts from patients with SSc and other fibrotic conditions have short primary cilia compared to fibroblasts from matched healthy controls [16]. We further characterised primary cilia length in dermal fibroblasts isolated from established dcSSc and Very Early Diagnosis Of Systemic Sclerosis (VEDOSS) patients. Compared to cells from healthy controls, SSc and VEDOSS fibroblasts were constitutively significantly shorter (Fig. 1A and 1B), a feature that was stably maintained over serial *in vitro* passaging. Primary cilia length was homogeneous across fibroblasts from different donors within each group (Fig. S1A) and followed a quasi-normal distribution, with a higher percentage of shorter primary cilia in SSc fibroblasts and a reduced overlap in cilia length with healthy controls (Fig. S1B). However, fibroblasts from healthy controls, SSc and VEDOSS patients had similar frequency (∼90%) of ciliation (Fig. S1C). Moreover, the short cilia phenotype was not an artifact of *ex vivo* culture since it was easily observed in primary cells within 2 passages from biopsy expansion (Fig. S1D), suggesting that hTERT-immortalised fibroblasts could be used for investigating the mechanisms underlying the ciliary phenotype.

**Figure 1.**
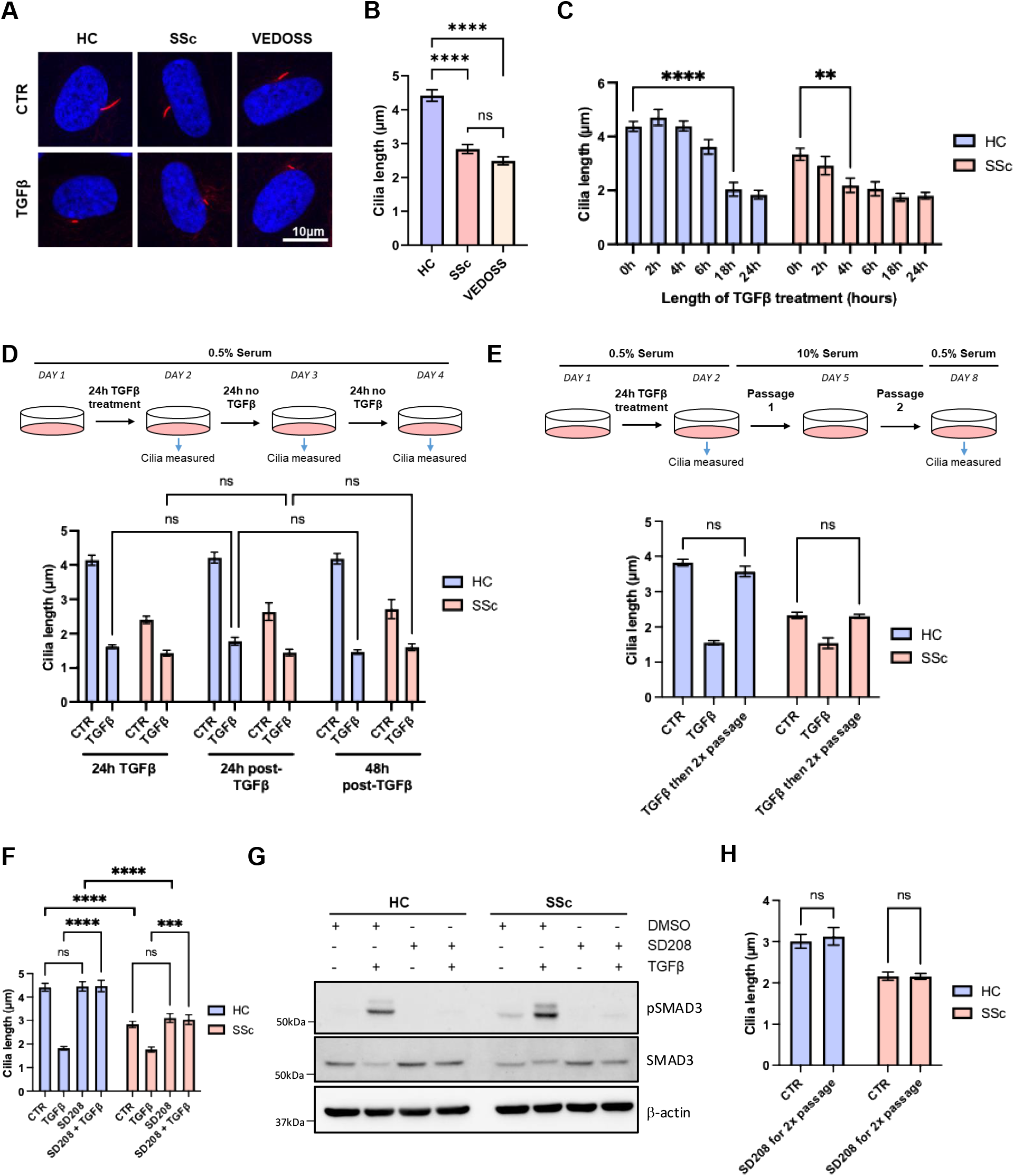
The short SSc cilia phenotype is stable and not driven by tonic TGFβ signalling. **(A)** Representative immunofluorescence images of primary cilia in healthy control (HC), systemic sclerosis (SSc), and VEDOSS (very early SSc) dermal fibroblasts under basal conditions (CTR) or following 24 h TGFβ stimulation. Acetylated α-tubulin marks the axoneme (red) and nuclei are stained with DAPI (blue). **(B)** Quantification of primary cilium length in HC, SSc, and VEDOSS fibroblasts under basal conditions. n=3 donors per group, a minimum of 100 cilia were quantified per sample. **(C)** Time course of TGFβ-induced cilia shortening in HC and SSc fibroblasts. n=3 donors per group, a minimum of 100 cilia were quantified per sample. **(D)** Quantification of cilium length in HC and SSc fibroblasts subjected to a 24 h TGFβ pulse followed by washout and recovery in 0.5% serum for 24 and 48 h. n=3 experimental repeats. **(E)** Quantification of cilium length in HC and SSc fibroblasts after sequential serum starvation, TGFβ exposure, and two passages in 10% serum. n=3 experimental repeats. **(F)** Cilium length in HC and SSc fibroblasts treated with the TGFβ receptor inhibitor SD208 in the presence or absence of exogenous TGFβ. n=3 experimental repeats. **(G)** Representative western blot of pSMAD3, total SMAD3, and βactin in HC and SSc fibroblasts treated with TGFβ and/or SD208. n=3 donors per group. **(H)** Quantification of cilium length in HC and SSc fibroblasts cultured in full growth medium in the presence of DMSO (CTR) or SD208 for 2 passages. n=3 experimental repeats. All data panels were analysed by ANOVA (ns, non-significant; ** P<0.01; *** P<0.001; ****P<0.0001).

Constitutive TGFβ signalling activation is a well-known feature in SSc fibroblasts [3]. In addition to its potent profibrotic effects, exogenous TGFβ1 treatment was shown to reduce the length of primary cilia in stromal cells including tenocytes, skin fibroblasts and lung fibroblasts [16, 17]. To better characterize this effect, healthy and SSc skin fibroblasts were allowed to form cilia and then incubated with 10 ng/ml TGFβ1. TGFβ1 induced a time-dependent reduction in cilia length in healthy fibroblasts, as well as an additional shortening in SSc fibroblasts, with statistically significant shortening occurring earlier (after only 4 h) in SSc cells compared with 18 h in healthy fibroblasts (Fig. 1C), while no changes in cilia frequency were detected (Fig. S1E). Moreover, TGFβ1 induced primary cilia shortening in all fibroblast types including immortalised fibroblasts (Fig. S1F). Next, we investigated if the effect of TGFβ1 on primary cilia length was reversible, for this purpose, fibroblasts were exposed to TGFβ1 for 24h, followed by fresh media without TGFβ1, and cilia length quantification at 24 h and 48 h post-withdrawal. In this experimental setting with low serum concentrations, in which the cell cycle is arrested at G1, cilia length reduction by TGFβ1 was maintained over 48 h (Fig. 1D). However, when cells were allowed to resume the cell cycle in fresh media containing 10% serum upon TGFβ1 removal, cilia length was re-established in both healthy and SSc cells after 48 h (2 passages), returning to their respective baseline levels (Fig. 1E).

To determine if the constitutive short cilia phenotype of SSc fibroblasts was mediated by canonical TGFβ signalling activation, inhibition of this signalling pathway was conducted. Blocking TGFBRI activation with SD208 prevented cilia shortening (Fig. 1F) and increased phospho-SMAD3 levels (Fig. 1G) in both healthy and SSc fibroblasts in response to TGFβ1, demonstrating the inhibitor’s efficacy; however, it did not restore cilia length in SSc fibroblasts to levels observed in healthy controls. Finally, we cultured SSc fibroblasts in the presence of SD208 and 10% serum for 2 passages to allow cilia disassembly and reformation during the cell cycle. Even under this condition, primary cilia on SSc fibroblasts remained significantly shorter than those of healthy control fibroblasts (Fig. 1H), suggesting that the constitutive short cilia phenotype is not the consequence of upstream canonical TGFβ signalling activation.

### Non-canonical TGFβ signalling through ROCK2 partially mediates its effect on cilia length

Next, we investigated the mechanistic basis of primary cilia shortening in fibrosis, both in basal condition in SSc and the additional shortening induced by TGFβ1 treatment. First, we tested whether canonical or non-canonical TGFβ signalling drives rapid cilia shortening induced by TGFβ1 treatment. To inhibit canonical Smad-mediated signalling, we downregulated SMAD3 expression in healthy fibroblasts by RNA silencing, followed by TGFβ1 treatment. A reduction of ∼70% of SMAD3 levels (Fig. 2A) did not prevent TGFβ-induced reduction of primary cilia length (Fig. 2B) nor affect the incidence of ciliation (Fig. S2A), in these fibroblasts. Interestingly, reducing cellular SMAD3 levels did not change the length of primary cilia on healthy control or SSc fibroblasts in resting conditions or after TGFβ1 treatment (Fig. 2B), suggesting that the mechanisms by which TGFβ1 induce primary cilia shortening might be non-canonical.

**Figure 2.**
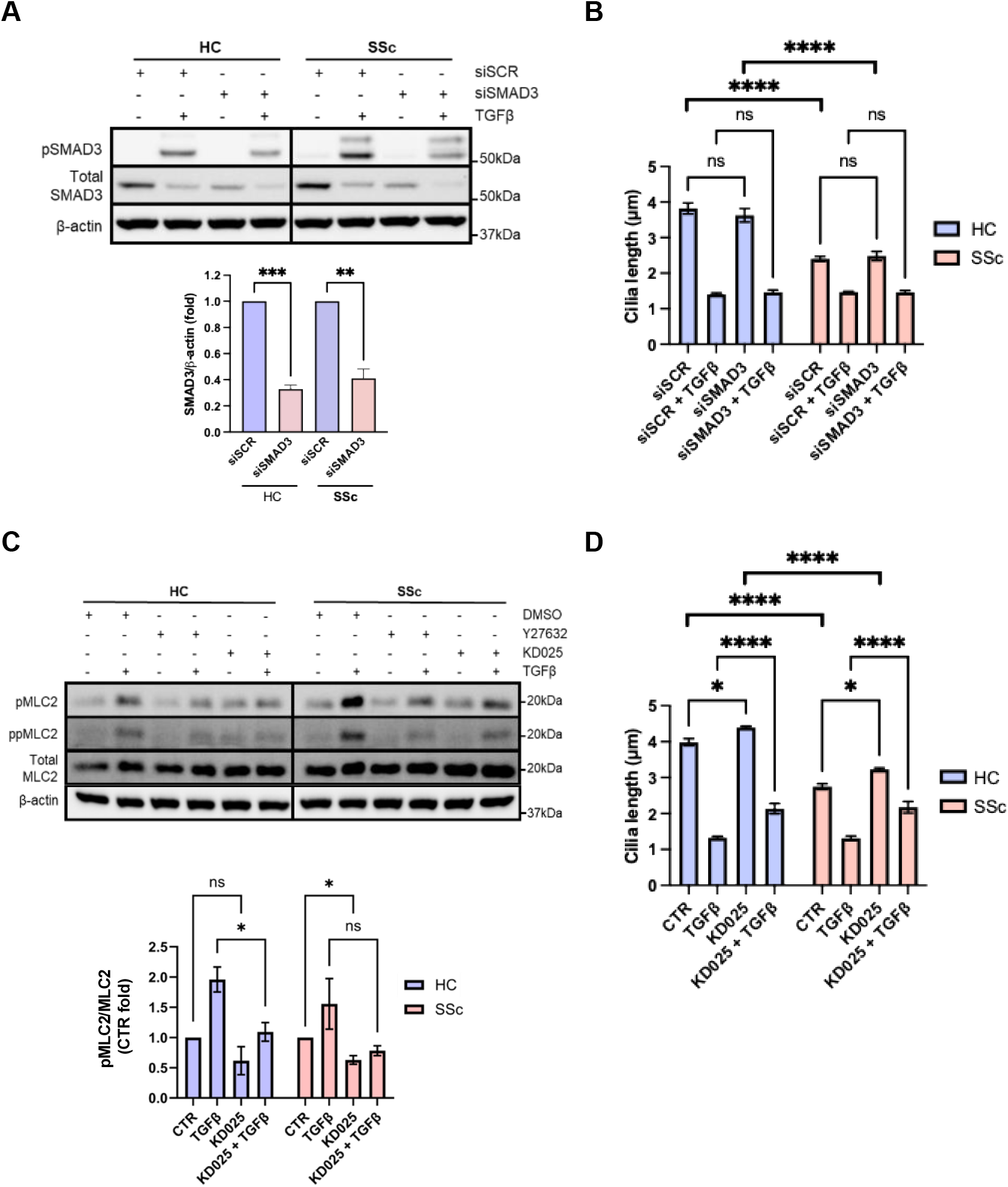
TGFβ-induced cilia shortening is mechanistically distinct from the basal short-cilia phenotype of SSc fibroblasts. **(A)** Top: representative western blot showing the effect of SMAD3 siRNA (siSMAD3) on total SMAD3 protein level and on TGFβ-induced SMAD3 phosphorylation in healthy control (HC) and systemic sclerosis (SSc) fibroblasts, compared to scrambled siRNA (siSCR). Bottom: Densitometric quantification of SMAD3 in control (siSCR) and siSMAD3-treated HC and SSc fibroblasts (n=3 individual donors per group). **(B)** Quantification of cilium length in HC and SSc fibroblasts treated with siSCR or siSMAD3 and stimulated for 24 h with TGFβ. n=3 individual donors per group. **(C)** Top: representative western blots showing pMLC2, ppMLC2, and total MLC2, in response to treatments TGFβ with or without KD025 in three HC donor and three SSc donor fibroblast lines. Bottom: Densitometric quantification of pMLC2 normalised to MLC2 in the same condition. **(D)** Cilium length in HC and SSc fibroblasts treated with the ROCK2 inhibitor KD205 ± TGFβ. Mean cilia length was measured in three HC donor and three SSc donor fibroblast lines treated as indicated. All data panels were analysed by ANOVA (ns, non-significant; * P<0.05; ****P<0.0001).

Multiple mechanisms of primary cilia resorption in response to diverse mitogenic stimuli have been reported. These include actin polymerisation downstream of Rho-dependent kinase (ROCK) [18]. Since TGFβ can stimulate ROCK activity by a non-canonical pathway independently of SMADs [19], we sought to determine the effect of ROCK on cilia length. There are two isoforms of ROCK and pan-ROCK inhibitors cause cell rounding and detachment. Therefore, we evaluated the effect of a ROCK2-specific inhibitor (KD205), which has more subtle effects on overall cellular health. Myosin light chain 2 (MLC2) phosphorylation, a surrogate of ROCK activity, was strongly increased by TGFβ1 in both healthy and SSc fibroblasts and partly prevented by KD205 (Fig. 2C). KD205 treatment induced a modest increase in cilia length in both healthy and SSc fibroblasts under basal conditions but did not abolish their relative length difference (Fig. 2D), and it did not change the percentage of ciliated cells (Fig. S2B). However, inhibition of ROCK2 blocked cilia shortening induced by TGFβ1 treatment (Fig. 2D), suggesting that TGFβ-induced cilia resorption is mediated by a RhoA/ROCK dependent non-canonical pathway.

### The short primary cilia phenotype and increased contractility of SSc fibroblasts involves augmented AURKA and HDAC6 activity

Another well studied mechanism underlying ciliary shortening is deacetylation of axonemal acetylated-α-tubulin by histone deacetylase 6 (HDAC6), which in turn promotes ciliary disassembly [13]. In agreement with our recent publication [16], treatment with the HDAC6 inhibitor Tubacin increased primary cilia length specifically in SSc fibroblasts, but not in healthy control cells (Fig. 3A and 3B), reducing their relative length difference to <25%. As expected, Tubacin treatment dose-dependently increased acetylated-α-tubulin levels in both healthy and SSc cell lines (Fig. 3C and Fig. S3).

**Figure 3.**
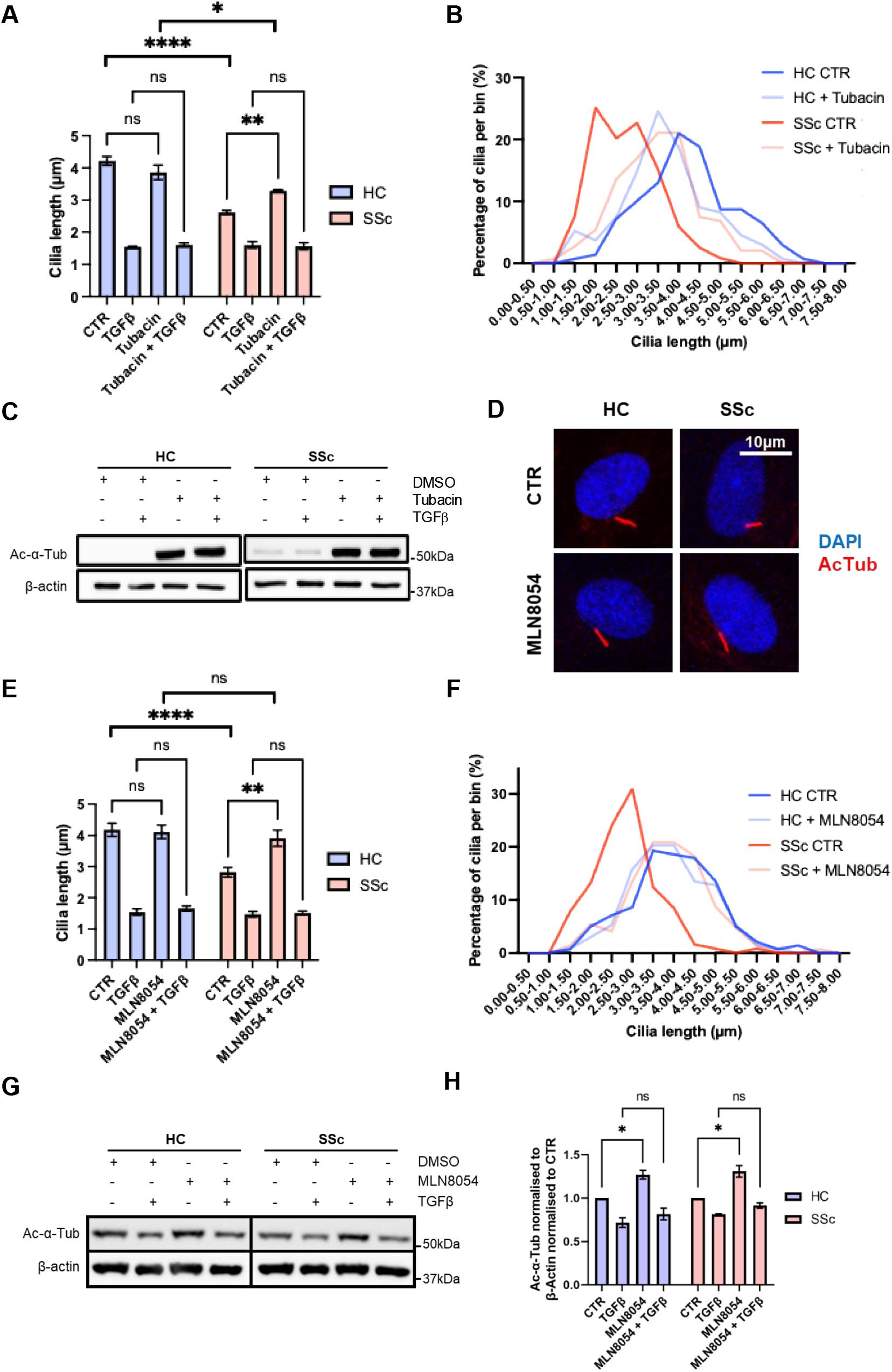
The AURKA/HDAC6 axis is required for maintenance of short cilia in SSc fibroblasts, but not for TGFβ-induced cilia resorption. **(A)** Quantification of primary cilium length in HC and SSc fibroblasts treated with vehicle or Tubacin and incubated with DMSO (CTR) or TGFβ for 24 h. n=3 donors per group. **(B)** Primary cilia length distribution in HC and SSc fibroblasts treated with vehicle or Tubacin. **(C)** Immunoblot of acetylated-α-tubulin levels in HC and SSc fibroblasts treated with vehicle or Tubacin and incubated with DMSO (CTR) or TGFβ for 24 h. **(D)** Representative immunofluorescence images of primary cilia in HC and SSc fibroblasts treated with DMSO (CTR) or the AURKA inhibitor MLN8054 for 24 h. Acetylated α-tubulin marks the axoneme (red) and nuclei are stained with DAPI (blue). **(E)** Quantification of cilium length in HC and SSc fibroblasts treated with vehicle or MLN8054 and incubated with DMSO (CTR) or TGFβ for 24 h. n=3 donors per group. **(F)** Primary cilia length distribution in HC and SSc fibroblasts treated with vehicle or MLN8054. **(G)** Immunoblot of acetylated-α-tubulin levels in HC and SSc fibroblasts treated with vehicle or MLN8054 and incubated with DMSO (CTR) or TGFβ for 24 h. **(H)** Densitometric quantification of acetylated-α-tubulin level normalised to β-actin. n=3 donors per group. All data panels were analysed by ANOVA (ns, non-significant; *P,0.05; ** P<0.01; ****P<0.0001).

A key mechanism of HDAC6 activation is through phosphorylation by Aurora A kinase (AURKA) [20]. Therefore, we used the AURKA inhibitor MLN8054 to investigate if AURKA was involved in promoting the short primary cilia phenotype of SSc fibroblasts. MLN8054 reduced the level of phosphorylated AURKA at mitotic spindle foci, a measure of its efficacy (Fig. S4A and S4B). However, it did not fully block AURKA at the concentration used, as it would block the cell cycle. Remarkably, MLN8054 treatment resulted in elongation of the primary cilia of SSc fibroblasts to the same level of those of healthy control cells (Fig. 3D and 3E), without affecting primary cilia length on healthy fibroblasts (Fig. 3F). Interestingly, while MLN8054 increased acetylated-α-tubulin levels, as expected due to activation of HDAC6, the increase was much smaller than that elicited with Tubacin (Fig. 3G-H), suggesting that AURKA might regulate primary cilia length by more than one mechanism. Furthermore, we showed in parallel experiments that TGFβ1-induced cilia shortening is independent of AURKA and HDAC6 activation (Fig. 3A and 3E). Altogether, our findings indicate that an AURKA-mediated mechanism participates in promoting the constitutive short cilia phenotype of SSc fibroblasts and the additional shortening mediated by exogenous TGFβı treatment is due to other mechanisms, including ROCK2 activation.

### Inhibition of AURKA or HDAC6 reduces myofibroblast markers and fibroblasts contractility

We have recently reported that elongation of primary cilia in SSc fibroblasts with Tubacin reduces expression of profibrotic markers [16]. Here, we tested whether the same is achieved by inhibition of the upstream regulatory kinase AURKA, a target of effective drugs in advanced clinical trials. Treatment of fibroblasts with MLN8054 at a concentration sufficient to induce cilia elongation reduced *ACTA2, CCN2* and *COL1A1* mRNA levels in SSc fibroblasts, but not *FN1* (Fig. 4A). In comparison, treatment with Tubacin strongly reduced all markers (Fig. 4A), even in healthy control fibroblasts. In agreement with a reduction of *ACTA2* expression, MLN8054 treatment abolished the increased contractility of SSc in collagen gel contraction assays (Fig. 4B). This effect was exclusively observed in SSc fibroblasts, since increased contractility of healthy control cells was insensitive to AURKA inhibition (Fig. 4B). Altogether, these data show a correlation of primary cilia length and myofibroblast activation in SSc, both co-regulated by AURKA/HDAC6.

**Figure 4.**
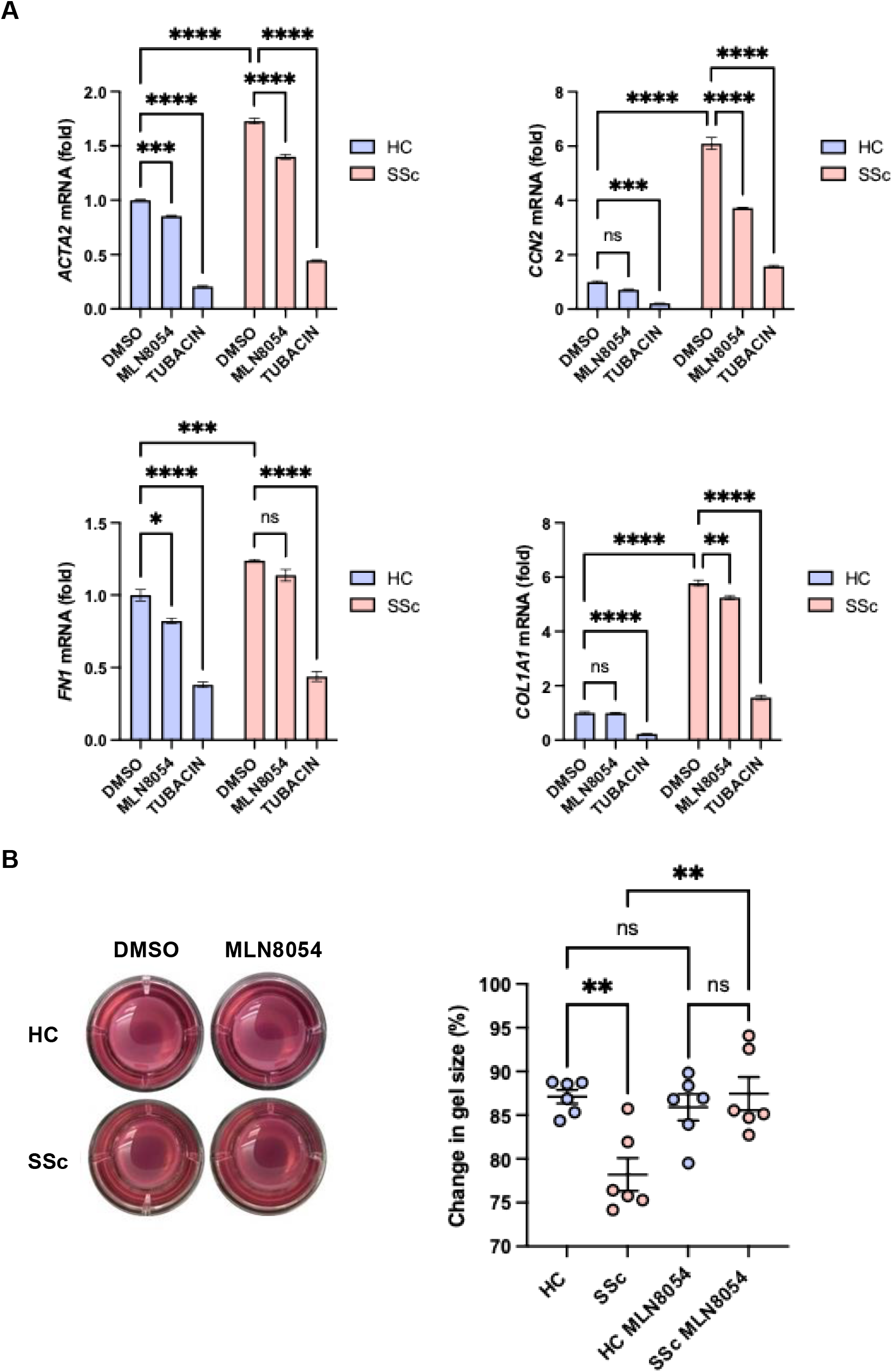
The AURKA/HDAC6 axis contributes to the pro-fibrotic phenotype of SSc fibroblasts. **(A)** Effect of MLN8054 and Tubacin compared to vehicle (DMSO) on the mRNA expression level of the indicated genes, as determined by qPCR using *GAPDH* as housekeeping control, in HC and SSc fibroblasts. n=3 donors per group. **(B)** Left: Representative images of collagen gels embedded with HC or SSc fibroblasts and treated with DMSO or MLN8054 in the presence of TGFβ for 48 h. Right: percentage of gel size change at 48 h compared to t=0 h by HC and SSc fibroblasts incubated with DMSO or MLN8054 in the presence of TGFβ. n= 6 experimental repeats. All data panels were analysed by ANOVA (ns, non-significant; *P,0.05; ** P<0.01; ***P<0.001; ****P<0.0001).

### Downregulation of Caveolin-1 reduces primary cilia length in healthy fibroblasts

Reduced Caveolin-1 (CAV1) levels are characteristic of SSc fibroblasts [21]. We have previously reported that silencing of CAV1 or blocking CAV1 function with a cell permeable peptide in normal fibroblasts increases pro-fibrotic markers expression, suggesting that reduction of CAV1 participates in SSc pathogenesis [21]. Here, we investigated if downregulation of CAV1 in normal fibroblasts from healthy controls is sufficient to trigger a reduction of primary cilia length like the one observed in SSc fibroblasts. Healthy fibroblasts were transduced with lentiviral particles encoding shRNA targeting CAV1 (shCAV1) or a scrambled sequence (shSCR). shCAV1 fibroblasts showed >70% reduction of CAV1 protein level compared to shSCR cells (Fig. 5A). Consistent with the observation that activated fibroblasts have shorter primary cilia, shCAV1 cells exhibited shorter primary cilia than shSCR cells, reaching a final length comparable to that of SSc fibroblasts and displaying a similar cilia length distribution (Fig. 5B, 5C, 5D and S5B), without disrupting cilia frequency (Fig. S5A). Given the similarity with SSc fibroblasts, we tested whether inhibiting AURKA could increase the cilia length of shCAV1 cells to control levels. As in SSc fibroblasts, MLN8054 treatment rescued the short cilia phenotype of shCAV1 cells, abolishing the differential cilia length to shSCR cells (Fig. 5E and 5F). Moreover, TGFβ1 treatment further decreased cilia length in both shSCR and shCAV1 fibroblasts to the same level through a mechanism independent of AURKA (Fig. 5G), without affecting cilia frequency (Fig. S5C). Thus, inhibition of AURKA can also rescue the short cilia phenotype in the shCAV1 model of SSc.

**Figure 5.**
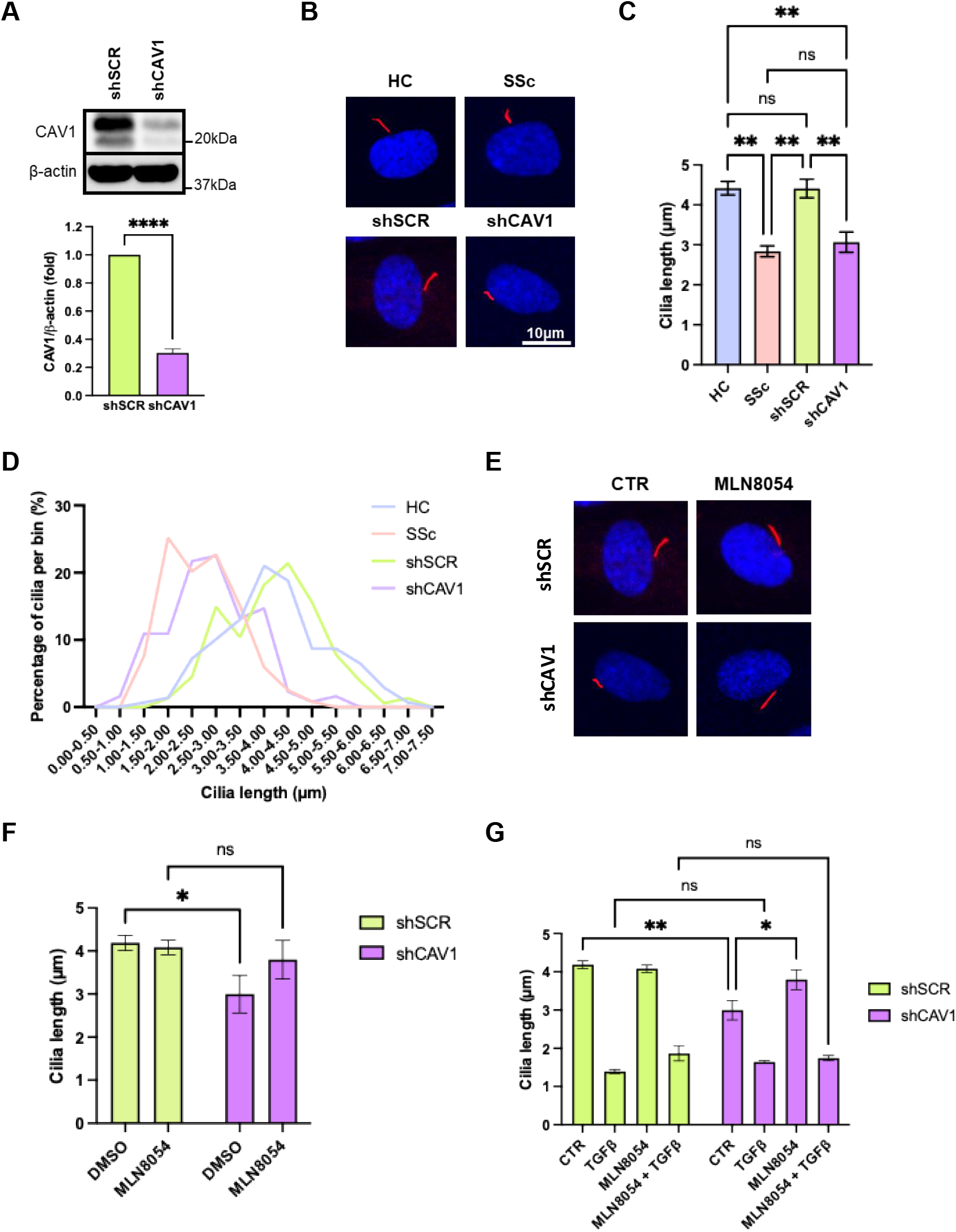
Downregulation of Caveolin-1 reduces primary cilium length, which is rescued by AURKA inhibition. **(A)** Protein levels of CAV1 in control (shSCR) and CAV1 silenced (shCAV1) healthy fibroblasts. Densitometric quantification of CAV1 levels normalised to β-actin. n=3 experimental repeats. **(B)** Representative immunofluorescence images of primary cilia (acetylated-α-tubulin, red; nucleus DAPI stain, blue) in shSCR and shCAV1 fibroblasts, compared to HC and SSc cells, in basal growing conditions. **(C)** Quantification of primary cilium length in shSCR and shCAV1 fibroblasts, compared to HC and SSc cells. n=3 experimental repeats, each measuring >100 primary cilia. **(D)** Primary cilia length distribution in shSCR and shCAV1 fibroblasts, compared to HC and SSc fibroblasts under basal conditions. **(E)** Representative immunofluorescence images of primary cilia in shSCR and shCAV1 fibroblasts incubated with DMSO (CTR) or the AURKA inhibitor MLN8054 for 24 h. Acetylated α-tubulin marks the axoneme (red) and nuclei are stained with DAPI (blue). **(F)** Quantification of cilium length in shSCR and shCAV1 fibroblasts treated with vehicle (DMSO) or MLN8054 for 24 h. n=3 experimental repeats. **(G)** Effect of MLN8054 or DMSO (CTR) on cilia length of shSCR and shCAV1 stimulated or not with TGFβ for 24 h. n=3 experimental repeats. All data panels were analysed by ANOVA (ns, non-significant; *P,0.05; ** P<0.01).

### Inhibition of AURKA or HDAC6 reduces fibroblast activation markers and contractility of shCAV1 fibroblasts

Since AURKA inhibition increases cilia length specifically in shCAV1 fibroblasts, we tested whether it would reduce their profibrotic phenotype. MLN8054 treatment partially reduced *ACTA2, CCN2, COL1A1* and *COL1A2* levels exclusively in shCAV1 but not in shSCR fibroblasts (Fig. 6A). When tested on a functional assay of fibroblasts contractility, MLN8054 also reduced gel contraction by shCAV1 cells, but not of shSCR cells, abrogating the profibrotic phenotype compared to shSCR (Fig. 6B). These findings suggests that a reduction of CAV1 expression in SSc could be upstream of aberrant AURKA activation in the process that leads to shortening of the primary cilia and cellular activation.

**Figure 6.**
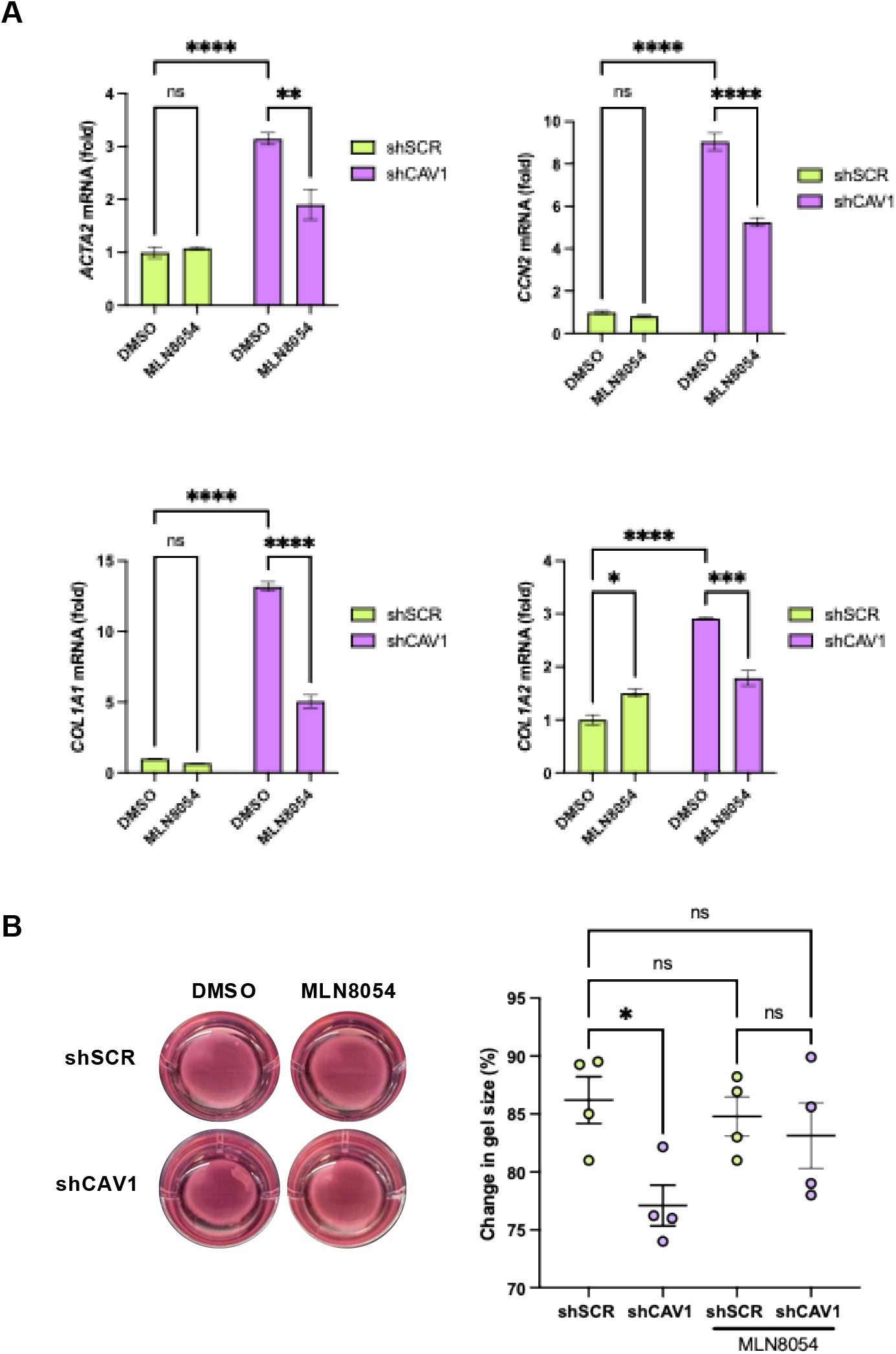
AURKA inhibition reduces the pro-fibrotic phenotype of shCAV1 fibroblasts. **(A)** Effect of MLN8054 compared to vehicle (DMSO) on the mRNA expression level of the indicated genes, as determined by qPCR using *GAPDH* as housekeeping control, in control (shSCR) and CAV1 silenced (shCAV1) fibroblasts. n=3 experimental repeats. **(B)** Left: Representative images of collagen gels embedded with shSCR or shCAV1 fibroblasts and treated with DMSO or MLN8054 in the presence of TGFβ for 48 h. Right: percentage of gel size change at 48 h compared to t=0 h by shSCR and shCAV1 fibroblasts incubated with DMSO or MLN8054 in the presence of TGFβ. n= 4 experimental repeats. All data panels were analysed by ANOVA (ns, non-significant; *P,0.05; ** P<0.01; ***P<0.001; ****P<0.0001).

## DISCUSSION

Our results demonstrate that dermal fibroblasts isolated from SSc patients have a stable short primary cilia phenotype that is evident before clinically detectable fibrosis, suggesting a potential role early in disease progression. Short cilia are evident in primary fibroblasts upon isolation and maintain their characteristics through culture passaging and upon immortalisation. In addition, our data indicate that the short cilia phenotype observed in SSc fibroblasts is not mediated by constitutive activation of canonical TGFβ signalling. However, we found that TGFβ1 induces cilia shortening in healthy control fibroblasts and further shortening in SSc fibroblasts to a similar final length in a reversible manner. Notably, this process is accelerated in SSc cells and is mediated, at least in part, by activation of Rho/ROCK signalling.

Our findings demonstrate that AURKA-mediated cilia disassembly mechanisms play a key role in maintaining the short cilia in SSc fibroblasts. An AURKA inhibitor increased SSc primary cilia length to the level of control cells without modifying cilia length of control cells, showing a selective effect on diseased fibroblasts. Inhibition of AURKA reduces the enhanced contractility displayed by SSc fibroblasts, which is accompanied by reduction in the expression of some profibrotic markers. AURKA is a central regulator of primary cilia disassembly through phosphorylation and activation of HDAC6, the α-tubulin deacetylase. In this study, HDAC6 inhibition with tubacin rescued cilia length in SSc cells and blocked profibrotic marker expression, in agreement with our previous report [16]. This observation suggests that the short ciliary phenotype is directly associated to fibroblast activation, highlighting the interdependence of cilia morphology and signalling modulation.

We and others have previously identified CAV1 as a critical regulator of fibrotic signalling pathways [22]. CAV1 expression is significantly reduced in the affected skin and lungs of SSc patients, contributing to enhanced TGFβ signalling with persistent Smad3 activation, and to excessive collagen production, hallmark features of SSc-associated fibrosis [21]. Genetic studies also suggest that *CAV1* polymorphisms may influence susceptibility and progression of SSc-related ILD [23]. In vitro studies using SSc fibroblasts and *Cav1* knockout mouse fibroblasts have demonstrated that loss of CAV1 is sufficient to induce a profibrotic phenotype, while restoration of CAV1 function via a cell-permeable scaffolding peptide reverses fibrosis and normalizes fibroblast behaviour [24]. Remarkably, we found that downregulation of CAV1 in dermal fibroblasts from healthy controls causes a reduction of primary cilia length indistinguishable from the SSc phenotype. In this cellular model of fibrosis, inhibition of AURKA or HDAC6 rescued cilia length and reduced the level of fibrotic markers and contractility, as seen in SSc fibroblasts. We propose that this finding is a strong indication that loss of CAV1 promotes abnormal AURKA activation that underlies partial cilia resorption in SSc.

AURKA activation is tightly controlled by multiple upstream signals. One well-characterized mechanism involves the interaction of AURKA with calcium-bound calmodulin (Ca^2+^/CaM), essential for recruitment and activation of AURKA at the basal body during interphase [25]. It is possible that CAV1 knockdown increases Ca^2+^//CaM in the cytosol, leading to increased activation of AURKA. Knockdown of CAV1 in human airway smooth muscle cells and in breast cancer cells has been shown to significantly reduce store-operated calcium entry (SOCE), increasing cytosolic Ca^2+^[26, 27]. This reduction was shown to be primarily due to decreased expression and membrane localization of Orai1 in airway smooth muscle cells. Therefore, it is possible that changes in Ca^2+^are upstream of AURKA activation and cilia shortening in SSc. Beyond Ca^2+^//CaM, proteins such as trichoplein keratin filament-binding protein (TCHP) and NEDD9/HEF1 have been implicated in stabilizing and activating AURKA at the ciliary base, contributing to its role in ciliary disassembly [28] [20]. Downregulation of SPAG17, a protein localised at the ciliary base, is another feature of SSc [29]. SPAG17 knockout cells are characterised by short cilia [30]; however, it is unknown if this is associated with concurrent downregulation of CAV1 and activation of AURKA. Future studies will be required to determine the trigger of the aberrant AURKA activation in SSc and the role of CAV1 and SPAG17 in the process.

Despite the association of CAV1 downregulation and primary cilia shortening described in this study, the opposite has been reported in epithelial cells with long primary cilia [18] In cells deficient of the Cav1α splicing isoform, apical RhoA activity is significantly reduced, which disrupts the organization of the actin cytoskeleton [18]. This disruption is accompanied by an accumulation of Rab11^+^ vesicles at the centrosome, suggesting altered vesicle trafficking dynamics. Further downstream, the RhoA effectors ROCK and DIA1 are implicated in this regulatory pathway, as their activity contributes to maintaining appropriate ciliary length [18]. It is important to note that these findings were observed in MDCK cells, which lack apical caveolae, indicating that Cav1α probably functions in a non-caveolar context to regulate cilia dynamics in those cells.

Our findings suggest that suppressing the exacerbated activation of the AURKA/HDAC6 axis could be of therapeutic value for the treatment and prevention of SSc-associated fibrosis. However, AURKA extraciliary functions in the regulation of the cell cycle make it a problematic target in a non-oncological disorder. Identification of specific regulators of AURKA ciliary function may, however, prove attractive therapeutic targets and warrant further investigation.

## MATERIALS AND METHODS

### Cell culture

Human dermal fibroblasts were isolated from full thickness forearm punch biopsies from healthy control (HC) donors (n=3) or from patients satisfying either the ACR-EULAR classification criteria for diffuse cutaneous SSc (dcSSc) [31] (n=3), or VEDOSS classification criteria [32](n=4) under ethics (reference number STRIKE 15/NE/0211), as previously described [33]. Patient linked cell line information are outlined in Table S1. HC fibroblasts were matched in donor age and sex. Primary fibroblasts experiments were performed within five passages. Primary lines were also immortalised by transduction with a retrovirus expressing human telomerase reverse transcriptase (hTERT), as previously described [34]. Cells were cultured in 1 g/L Glucose Dulbecco’s Modified Eagle Medium (DMEM) (Thermo Fisher) with 10 % FBS (Thermo Fisher), 1 % Penicillin/Streptomycin (Merck), and 250 µg/mL Geneticin (Thermo Fisher) as selection for hTERT transduction.

Geneticin was excluded from culture medium while performing experiments.

### Gene silencing

To study the effect of Caveolin-1 (CAV1) downregulation, healthy dermal fibroblasts were transduced with short hairpin RNA (shRNA) GIPZ lentivirus systems (Open Biosystems) containing either scramble control shRNA (RHS4348) or CAV1 (V3LHS_312897-100991154) sequence specific shRNA. Constructs were introduced to fibroblasts and incubated for 48 h before selection for successful transduction with Puromycin (1 mg/mL, Life Technologies) for 10 days, as previously described [35]. During culture, selection was maintained with 0.5 μg/mL Puromycin, and continuation of knockdown monitored by Western Blot for CAV1. To inhibit the canonical TGFβ pathway, a pool of four FlexiTube GeneSolution siRNA (Qiagen) oligos specific to SMAD3 were used for post-transcriptional gene silencing. The four target sequences were as follows: AGCCTATACTTTGGCAGGTTA, AAGGAGCACCTTGACAGACTT, AAGAGATTCGAATGACGGTAA, and ATCAAGGGATTTCCTATGGAA. Oligos were combined to a total concentration of 40 nM and transfected into fibroblasts at 70% confluence with Lipofectamine 2000 (1 μg/mL, Thermo Fisher) before incubation for 72 h, including any additional treatments, before protein harvest or fixation for immunofluorescence. As a negative control, fibroblasts were transfected with a scrambled (SCR) siRNA oligo (Qiagen) with the following sequence: AATTCTCCGAACGTGTCACGT. Knockdown was monitored by Western Blot for SMAD3.

### Inhibitor treatments

All experimental treatments were administered after fibroblasts reached confluence and were subjected to 24 h of serum starvation in DMEM (Thermo Fisher) with 0.5 % FBS (Thermo Fisher) and 1 % Penicillin/Streptomycin (Merck). Ligands or small molecule inhibitors were then added to fibroblasts at their appropriate concentrations in serum starved media for 24 h or 48 h, as described. Fibroblasts were treated with recombinant TGFβ1 ligand (10 ng/mL, Merck), TGFβRI inhibitor SD208 (1 μM, Selleckchem), ROCK2 inhibitor KD025 (5 μM, Cayman Chemical), HDAC6 inhibitor Tubacin (10 μM, Sigma), and AURKA inhibitor MLN8054 (10 μM, APExBIO). For controls, fibroblasts were treated with an equal volume of vehicle (DMSO, Sigma). In cases where cells were treated with both an inhibitor and TGFβ, inhibitor treatment was introduced two hours before stimulation with TGFβ.

### Immunofluorescence

Cells were seeded in Nunc™ Lab-Tek™ II Chamber Slides (Thermo Fisher) and subjected to experimental treatments. Cells were fixed with 4 % paraformaldehyde in PBS for 15 minutes at room temperature. For immunofluorescence analysis, briefly cells were permeabilised with 0.2 % Triton X-100 (Sigma) in PBS for 15 minutes, 1 % BSA (Sigma) in PBS blocking solution for 10 minutes at room temperature, and then incubated at 4 [C with primary antibodies (Acetylated-α-Tubulin; Cell Signaling, Ki67; Proteintech, pAURKA (Thr288); Abcam) in 1 % BSA, followed by incubation with secondary antibody (Anti-Rabbit (Alexa Fluor 594) or Anti-Rabbit (Alexa Fluor 594), Invitrogen) for 2 hours at room temperature, protected from light. Cells were washed with 1 X PBS between all incubations. Coverslips were mounted to slides using Prolong Anti-Fade Mounting Medium with DAPI (Life Technologies). Immunofluorescence was visualised and images acquired with an LSM880 upright laser scanning confocal microscope (Zeiss) with ZEN microscopy software, using a 40X magnification Plan-Apochromat DIC oil immersion objective lens (Zeiss). Images were taken at 0.2 µm intervals through the entire depth of the cell monolayer, then combined into Z-stacks. Quantitative analysis of microscopy images for the measurement of primary cilia length and incidence was conducted using ImageJ software and manual analysis of a minimum of 100 individual cells from a minimum of three randomly selected fields of view per experimental replicate.

### Western blotting

Total proteins were extracted from fibroblasts in ice cold protein lysis buffer comprising of RIPA Buffer (Merck) with 1 X Protease Inhibitor (Merck) and 1 X Phosphatase Inhibitor (Merck). Protein concentration was determined by colorimetric assay using the Pierce Bicinchoninic Acid (BCA) Protein Assay kit (Thermo Fisher) as per the manufacturer’s instructions and resolved by sodium dodecyl sulfate– polyacrylamide gel electrophoresis (10%–15% Tris-Glycine). Proteins were transferred onto Hybond nitrocellulose membranes (Amersham) and probed with antibodies specific for αSMA (Abcam ab7817), β-Actin (Sigma A5441), CAV1 (Santa Cruz sc894), p53 (Santa Cruz sc126), acetylated-α-Tubulin (Cell Signaling 5335), MLC2 (Cell Signaling 8505), pMLC2 (Ser19) (Cell Signaling 3674), ppMLC2 (Ser19, Thr18) (Cell Signaling 3674), SMAD3 (Cell Signalling 9523), pSMAD3 (Abcam ab52903). Immunoblots were visualized with species-specific horseradish peroxidase (HRP)-conjugated secondary antibodies (Cell Signaling 7074 and 7076) and enhanced chemiluminescence (Thermo Scientific/Pierce) on a Bio-Rad ChemiDoc imaging system.

### Quantitative reverse transcription–polymerase chain reaction

RNA was extracted from cells using commercial RNA extraction kits (Zymo Research). RNA (1 μg) was reverse transcribed using complementary DNA synthesis kits (ThermoFisher Scientific). Quantitative reverse transcription– polymerase chain reactions (qRT-PCRs) were performed using SyBr Green PCR kits on a Thermocycler with primers specific for *ACTA2* (FWD 5’-TGT ATG TGG CTA TCC AGG CG-3’ and REV 5’-AGA GTC CAG CAC GAT GCC AG-3’), *CCN2* (GTG TGC ACT GCC AAA GAT GGT and TTG GAA GGA CTC ACC GCT G), *FN1* (ACA ACA CCG AGG TGA CTG AGA C and GGA CAC ACC GAT GCT TCC TGA G), *COL1A1* (CCT CCA GGG CTC CAA CGA G and TCT ATC ACT GTC TTG CCC CA), *COL1A2* (GAT GTT GAA CTT GTT GCT GAG C and TCT TTC CCC ATT CAT TTG TCT T), *GLI1* (GGA CCT GCA GAC GGT TAT CC and AGC CTC CTG GAG ATG TGC AT) and *GLI2* (AGC AGC AGC AAC TGT C and GAA TGG CGA CAG GGT TGA C). Data were analysed using the 2^-ΔΔCt^ method using the basal expression in healthy control fibroblasts as control. *GAPDH* served as a housekeeping gene (ACC CAC TCC TCC ACC TTT GA and CTG TTG CTG TAG CCA AAT TCG T).

### Cell contraction assays

Cell Contraction Assay kit (Cell Biolabs) was used as per the manufacturer’s instructions. Briefly, 200,000 fibroblasts per condition were resuspended in a collagen solution, and the subsequent cell-collagen gel mixture was allowed to polymerise in a 24-well cell culture plate at 37 °C for 1 h in the absence of CO2. DMEM with 0.5 % FBS, 1 % Penicillin/Streptomycin and TGFβ ligand (10 ng/mL) containing the appropriate treatments was then added on top of the polymerised gels before incubation for 16 h at 37 °C with 5 % CO2. The edges of the cell-collagen gels were then released from the wells using a sterile scalpel, and images of each gel were taken at 0 h, 1 h, 2 h, 4 h, 6 h, 8 h, 24 h, and 48 h post-release. Images were analysed using ImageJ software to measure the area of each gel at each time point and to calculate the percentage of gel contraction relative to the 0h time point area.

### Statistical analysis

Results presented as continuous variables are reported as mean ± standard error or mean ± standard deviation, as noted in figure legends. Categorical variables are reported as percentages. Tests used to determine statistical significance are described in figure legends, with significance shown on data graphs as * for P<0.05, ** for P<0.01, *** for P<0.001, and **** for P<0.0001. All statistical analyses were carried out in GraphPad Prism software (version 10.3.1 (464)).

## DECLARATIONS

### Ethics approval and consent to participate

This study was conducted as part of the observational study STRIKE, approved by NorthEast-Newcastle & North Tyneside 2 Research Ethics Committee (IRAS ID: 178638; REC Reference: 15/NE/0211). All participants voluntarily agreed to participate providing written informed consent.

### Consent for publication

Not applicable.

## Acknowledgements

We are indebted to Prof. Colin Johnson (University of Leeds) for useful discussions and provision of KD205.

## Authors’ contributions

RW: experimentation, data analysis, manuscript draft; BCR: experimentation, data analysis; PM: experimentation, data analysis; AJT: experimentation, data analysis; MET: conceptualisation, draft review, JV: conceptualisation, draft review, FDG: conceptualisation, draft review, funding acquisition; RLR: experimental design, supervision, manuscript draft; NRDG: conceptualisation, experimental design, supervision, manuscript draft, funding acquisition.

## REFERENCES

1. Lescoat, A., et al., Systemic sclerosis: pathogenic mechanisms and their implications for treatment. Semin Immunopathol, 2025. 47(1): p. 39.

2. Elhai, M., et al., Mapping and predicting mortality from systemic sclerosis. Ann Rheum Dis, 2017. 76(11): p. 1897–1905.

3. Younesi, F.S., et al., Fibroblast and myofibroblast activation in normal tissue repair and fibrosis. Nat Rev Mol Cell Biol, 2024. 25(8): p. 617–638.

4. Beyer, C. and J.H. Distler, Morphogen pathways in systemic sclerosis. Curr Rheumatol Rep, 2013. 15(1): p. 299.

5. Clement, C.A., et al., TGF-beta signaling is associated with endocytosis at the pocket region of the primary cilium. Cell Rep, 2013. 3(6): p. 1806–14.

6. Ho, E.K. and T. Stearns, Hedgehog signaling and the primary cilium: implications for spatial and temporal constraints on signaling. Development, 2021. 148(9).

7. Lancaster, M.A., J. Schroth, and J.G. Gleeson, Subcellular spatial regulation of canonical Wnt signalling at the primary cilium. Nat Cell Biol, 2011. 13(6): p. 700–7.

8. Schneider, L., et al., PDGFRalphaalpha signaling is regulated through the primary cilium in fibroblasts. Curr Biol, 2005. 15(20): p. 1861–6.

9. Wachten, D. and S.T. Christensen, Primary cilia signalling at a glance. J Cell Sci, 2025. 138(20).

10. Wang, L. and B.D. Dynlacht, The regulation of cilium assembly and disassembly in development and disease. Development, 2018. 145(18).

11. Kim, S., et al., Nek2 activation of Kif24 ensures cilium disassembly during the cell cycle. Nat Commun, 2015. 6: p. 8087.

12. Ishikawa, H. and W.F. Marshall, Ciliogenesis: building the cell’s antenna. Nat Rev Mol Cell Biol, 2011. 12(4): p. 222–34.

13. Ott, C.M. and S. Mukhopadhyay, Taking Down the Primary Cilium: Pathways for Disassembly in Differentiating Cells. Bioessays, 2025. 47(11): p. e70060.

14. Teves, M.E., et al., The Primary Cilium: Emerging Role as a Key Player in Fibrosis. Curr Rheumatol Rep, 2019. 21(6): p. 29.

15. Bhattacharyya, D., M.E. Teves, and J. Varga, The dynamic organelle primary cilia: emerging roles in organ fibrosis. Curr Opin Rheumatol, 2021. 33(6): p. 495–504.

16. Verma, P., et al., Morphological reprogramming of primary cilia length mitigates the fibrotic phenotype in fibroblasts across diverse fibrotic conditions. J Cell Sci, 2025.

17. Rowson, D.T., et al., Mechanical loading induces primary cilia disassembly in tendon cells via TGFbeta and HDAC6. Sci Rep, 2018. 8(1): p. 11107.

18. Rangel, L., et al., Caveolin-1alpha regulates primary cilium length by controlling RhoA GTPase activity. Sci Rep, 2019. 9(1): p. 1116.

19. Bhowmick, N.A., et al., Transforming growth factor-beta1 mediates epithelial to mesenchymal transdifferentiation through a RhoA-dependent mechanism. Mol Biol Cell, 2001. 12(1): p. 27–36.

20. Pugacheva, E.N., et al., HEF1-dependent Aurora A activation induces disassembly of the primary cilium. Cell, 2007. 129(7): p. 1351–63.

21. Del Galdo, F., et al., Decreased expression of caveolin 1 in patients with systemic sclerosis: crucial role in the pathogenesis of tissue fibrosis. Arthritis Rheum, 2008. 58(9): p. 2854–65.

22. Del Galdo, F., M.P. Lisanti, and S.A. Jimenez, Caveolin-1, transforming growth factor-beta receptor internalization, and the pathogenesis of systemic sclerosis. Curr Opin Rheumatol, 2008. 20(6): p. 713–9.

23. Manetti, M., et al., Evidence for caveolin-1 as a new susceptibility gene regulating tissue fibrosis in systemic sclerosis. Ann Rheum Dis, 2012. 71(6): p. 1034–41.

24. Tourkina, E., et al., Altered monocyte and fibrocyte phenotype and function in scleroderma interstitial lung disease: reversal by caveolin-1 scaffolding domain peptide. Fibrogenesis Tissue Repair, 2011. 4(1): p. 15.

25. Plotnikova, O.V., et al., Calmodulin activation of Aurora-A kinase (AURKA) is required during ciliary disassembly and in mitosis. Mol Biol Cell, 2012. 23(14): p. 2658–70.

26. Sathish, V., et al., Caveolin-1 regulation of store-operated Ca(2+) influx in human airway smooth muscle. Eur Respir J, 2012. 40(2): p. 470–8.

27. Zhu, H., et al., Caveolae/caveolin-1 are important modulators of store-operated calcium entry in Hs578/T breast cancer cells. J Pharmacol Sci, 2008. 106(2): p. 287–94.

28. Inoko, A., et al., Trichoplein and Aurora A block aberrant primary cilia assembly in proliferating cells. J Cell Biol, 2012. 197(3): p. 391–405.

29. Sapao, P., et al., Reduced SPAG17 Expression in Systemic Sclerosis Triggers Myofibroblast Transition and Drives Fibrosis. J Invest Dermatol, 2023. 143(2): p. 284–293.

30. Teves, M.E., et al., Spag17 deficiency results in skeletal malformations and bone abnormalities. PLoS One, 2015. 10(5): p. e0125936.

31. van den Hoogen, F., et al., 2013 classification criteria for systemic sclerosis: an American college of rheumatology/European league against rheumatism collaborative initiative. Ann Rheum Dis, 2013. 72(11): p. 1747–55.

32. Avouac, J., et al., Preliminary criteria for the very early diagnosis of systemic sclerosis: results of a Delphi Consensus Study from EULAR Scleroderma Trials and Research Group. Ann Rheum Dis, 2011. 70(3): p. 476–81.

33. Ross, R.L., et al., Biological hallmarks of systemic sclerosis are present in the skin and serum of patients with Very Early Diagnosis of Systemic Sclerosis (VEDOSS). Rheumatology (Oxford), 2025. 64(6): p. 3606–3617.

34. Gillespie, J., et al., Transforming Growth Factor beta Activation Primes Canonical Wnt Signaling Through Down-Regulation of Axin-2. Arthritis Rheumatol, 2018. 70(6): p. 932–942.

35. Liakouli, V., et al., Scleroderma fibroblasts suppress angiogenesis via TGF-beta/caveolin-1 dependent secretion of pigment epithelium-derived factor. Ann Rheum Dis, 2018. 77(3): p. 431–440.

