## Supplementary figures for "Aurora A kinase activation contributes to the fibrotic phenotype in Systemic Sclerosis through primary cilia shortening"

### Slide 1
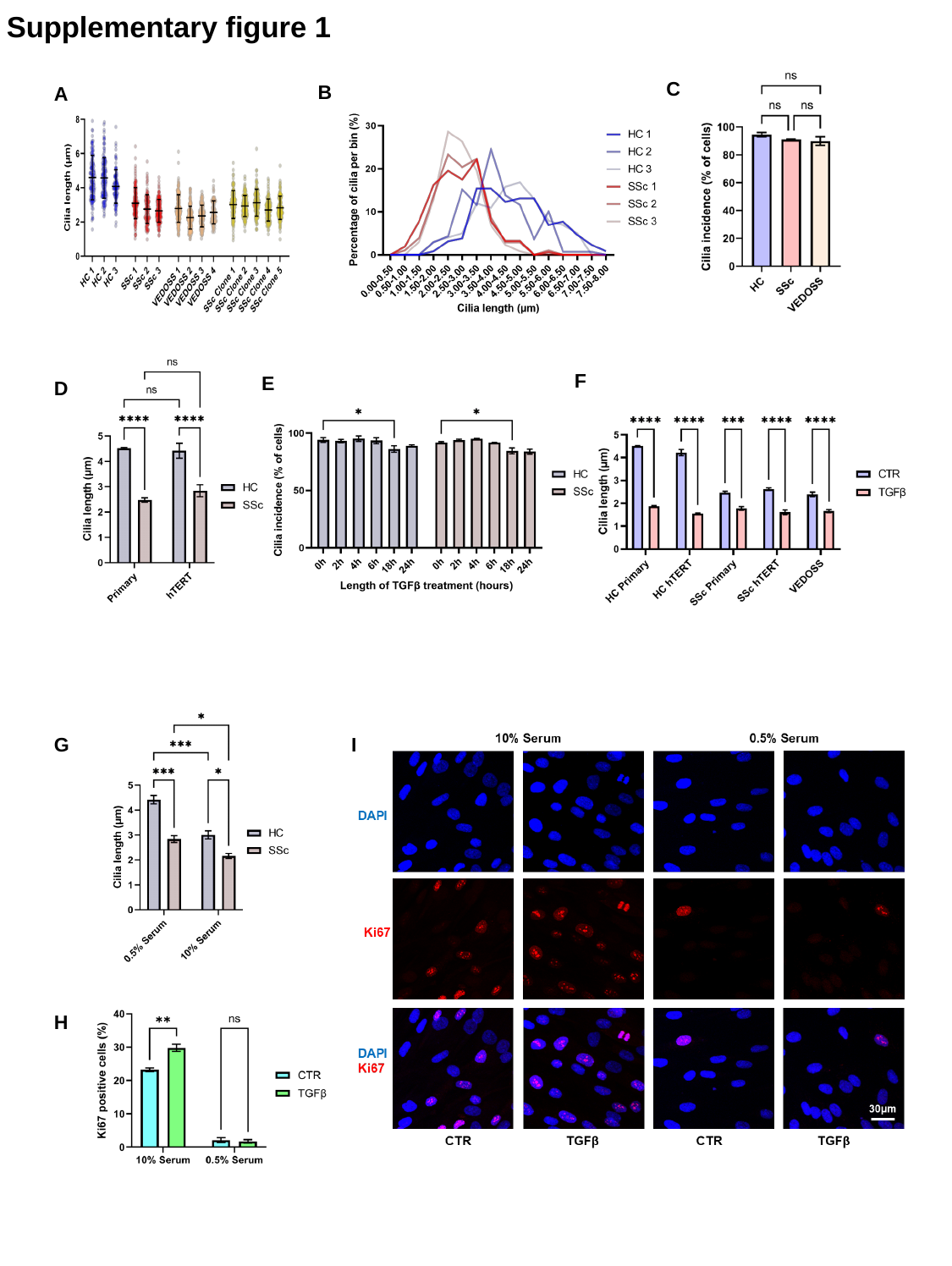

Supplementary figure 1
C
B
A
F
E
D
G
I
H

### Slide 2
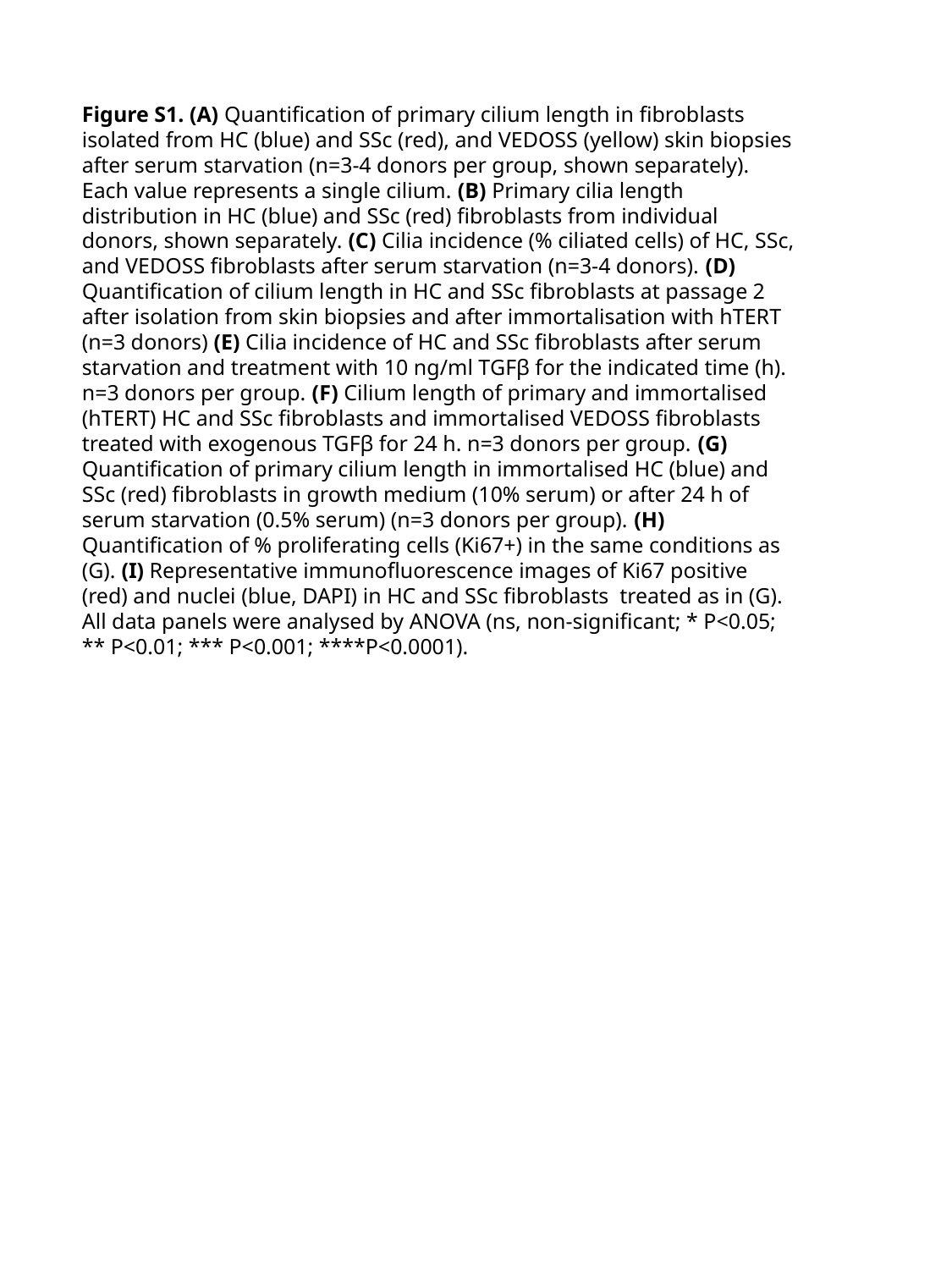

Figure S1. (A) Quantification of primary cilium length in fibroblasts isolated from HC (blue) and SSc (red), and VEDOSS (yellow) skin biopsies after serum starvation (n=3-4 donors per group, shown separately). Each value represents a single cilium. (B) Primary cilia length distribution in HC (blue) and SSc (red) fibroblasts from individual donors, shown separately. (C) Cilia incidence (% ciliated cells) of HC, SSc, and VEDOSS fibroblasts after serum starvation (n=3-4 donors). (D) Quantification of cilium length in HC and SSc fibroblasts at passage 2 after isolation from skin biopsies and after immortalisation with hTERT (n=3 donors) (E) Cilia incidence of HC and SSc fibroblasts after serum starvation and treatment with 10 ng/ml TGFβ for the indicated time (h). n=3 donors per group. (F) Cilium length of primary and immortalised (hTERT) HC and SSc fibroblasts and immortalised VEDOSS fibroblasts treated with exogenous TGFβ for 24 h. n=3 donors per group. (G) Quantification of primary cilium length in immortalised HC (blue) and SSc (red) fibroblasts in growth medium (10% serum) or after 24 h of serum starvation (0.5% serum) (n=3 donors per group). (H) Quantification of % proliferating cells (Ki67+) in the same conditions as (G). (I) Representative immunofluorescence images of Ki67 positive (red) and nuclei (blue, DAPI) in HC and SSc fibroblasts treated as in (G). All data panels were analysed by ANOVA (ns, non-significant; * P<0.05; ** P<0.01; *** P<0.001; ****P<0.0001).

### Slide 3
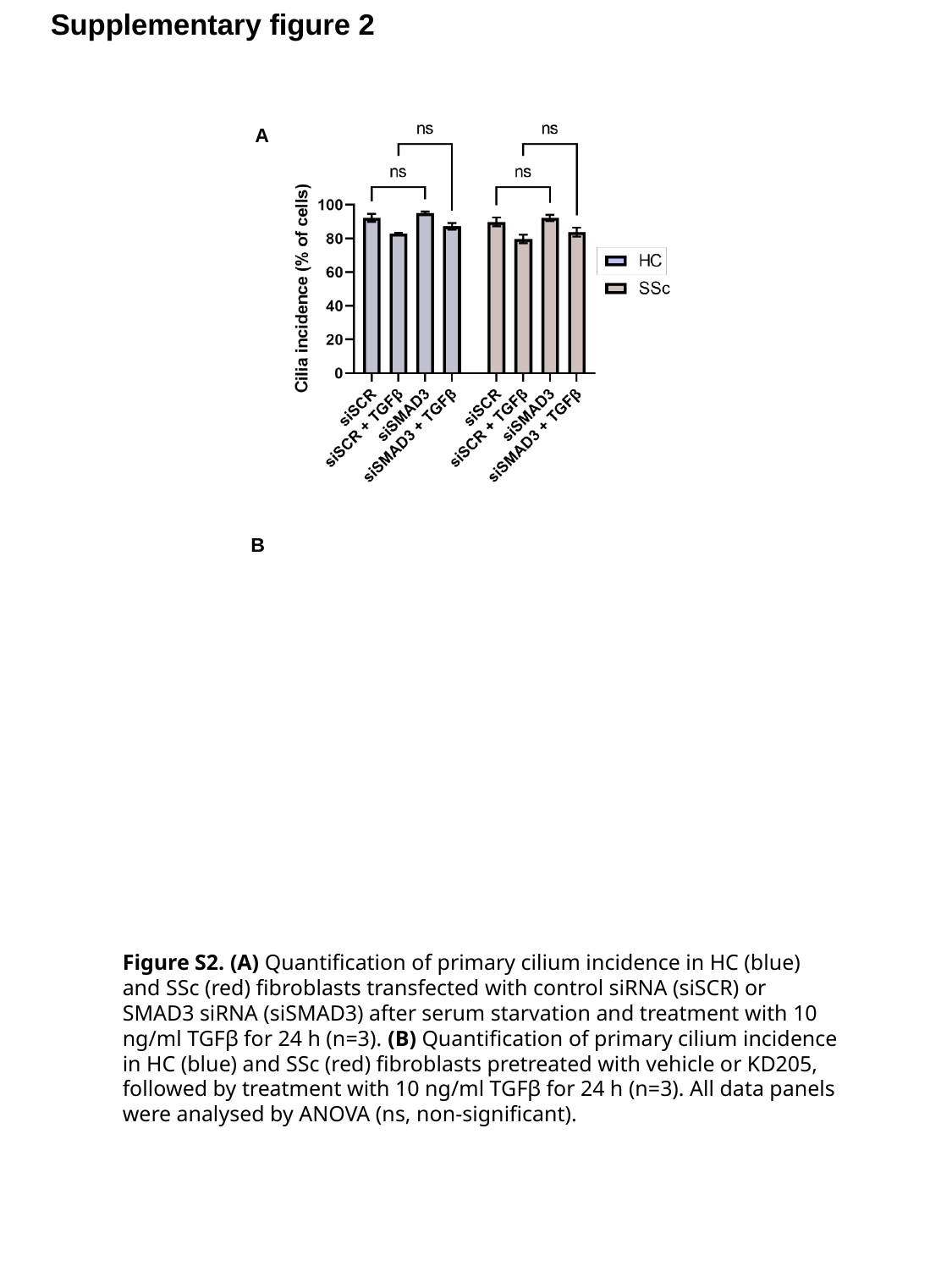

Supplementary figure 2
A
B
Figure S2. (A) Quantification of primary cilium incidence in HC (blue) and SSc (red) fibroblasts transfected with control siRNA (siSCR) or SMAD3 siRNA (siSMAD3) after serum starvation and treatment with 10 ng/ml TGFβ for 24 h (n=3). (B) Quantification of primary cilium incidence in HC (blue) and SSc (red) fibroblasts pretreated with vehicle or KD205, followed by treatment with 10 ng/ml TGFβ for 24 h (n=3). All data panels were analysed by ANOVA (ns, non-significant).

### Slide 4
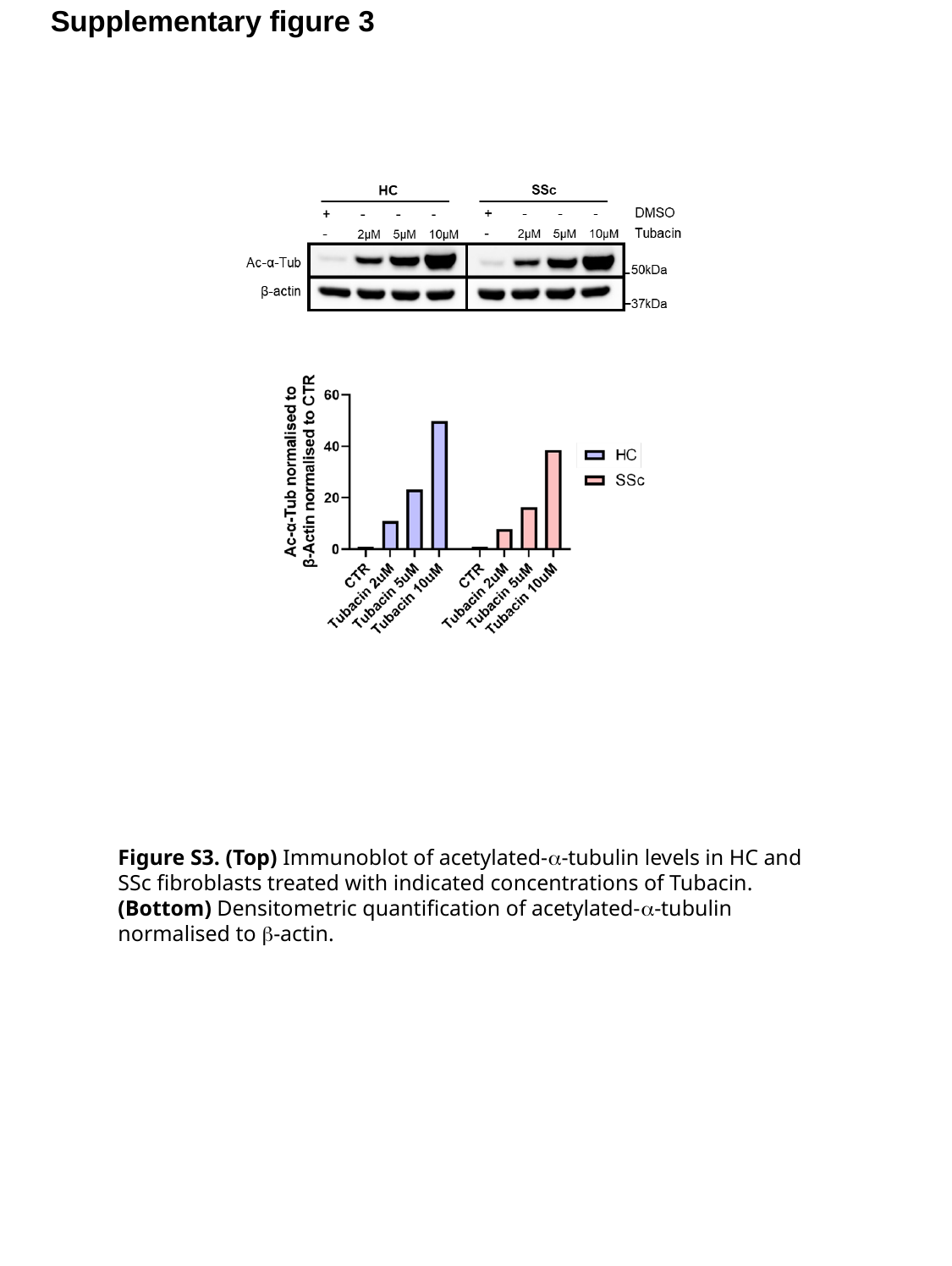

Supplementary figure 3
Figure S3. (Top) Immunoblot of acetylated-α-tubulin levels in HC and SSc fibroblasts treated with indicated concentrations of Tubacin. (Bottom) Densitometric quantification of acetylated-α-tubulin normalised to β-actin.

### Slide 5
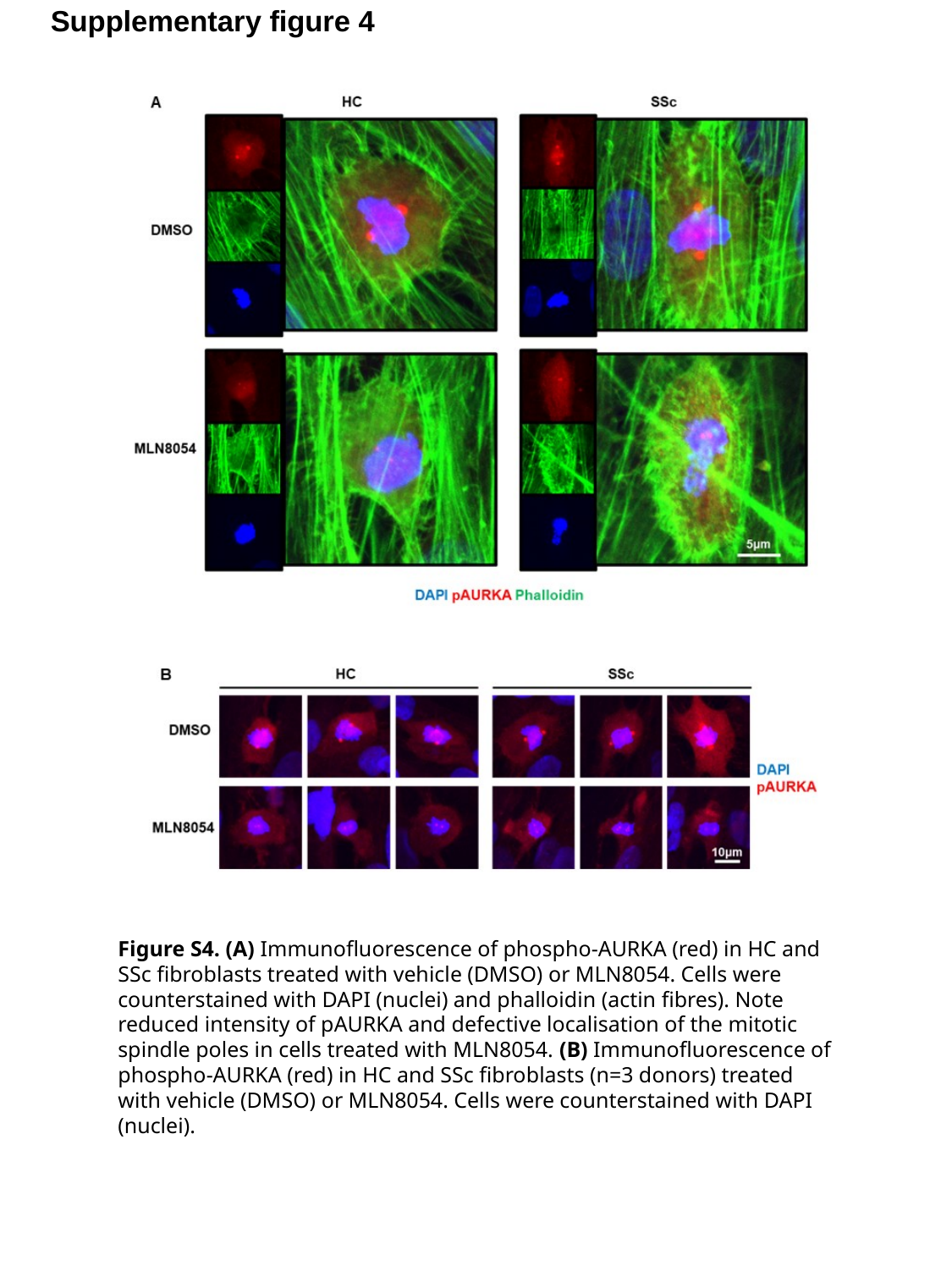

Supplementary figure 4
Figure S4. (A) Immunofluorescence of phospho-AURKA (red) in HC and SSc fibroblasts treated with vehicle (DMSO) or MLN8054. Cells were counterstained with DAPI (nuclei) and phalloidin (actin fibres). Note reduced intensity of pAURKA and defective localisation of the mitotic spindle poles in cells treated with MLN8054. (B) Immunofluorescence of phospho-AURKA (red) in HC and SSc fibroblasts (n=3 donors) treated with vehicle (DMSO) or MLN8054. Cells were counterstained with DAPI (nuclei).

### Slide 6
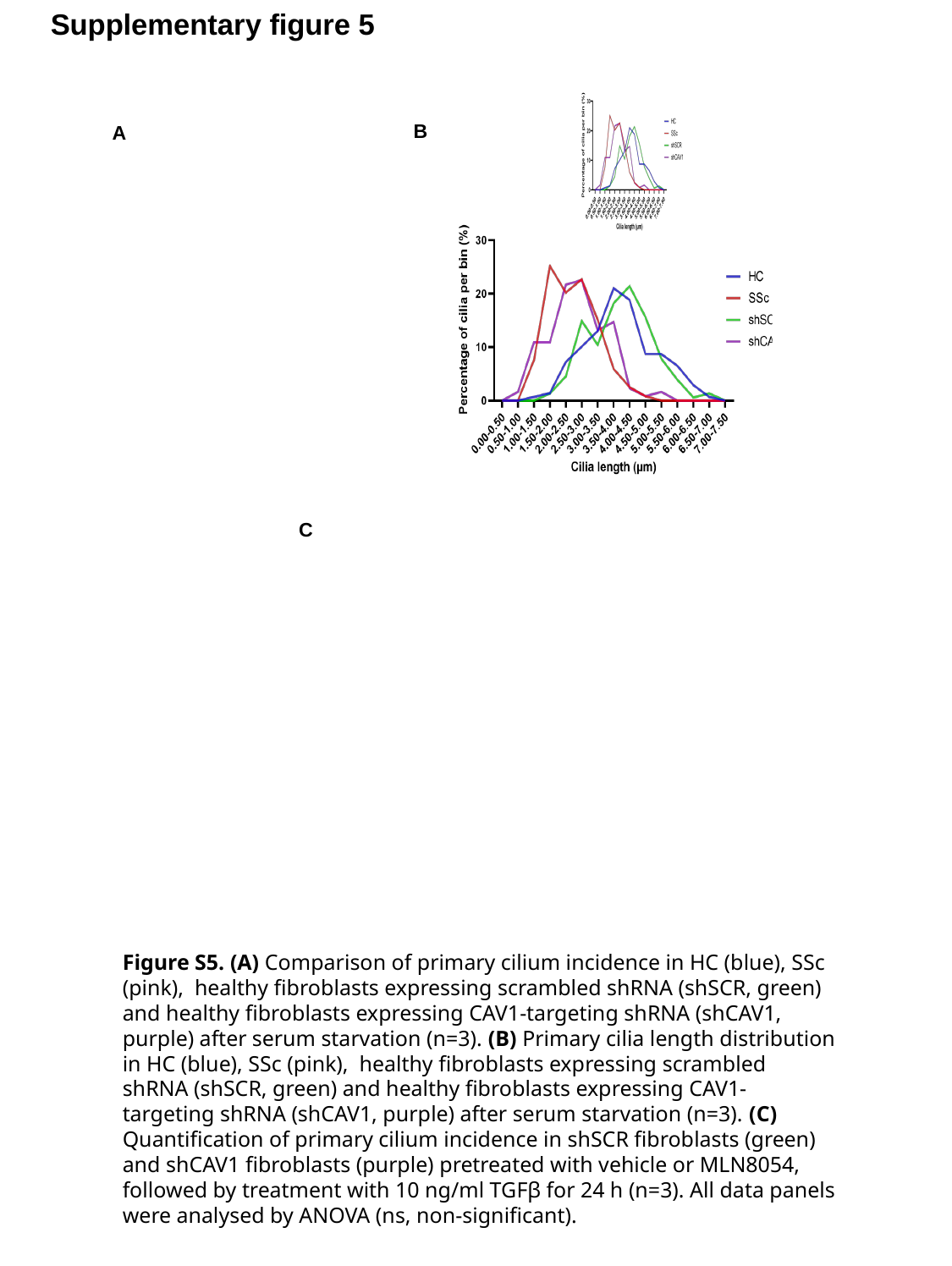

Supplementary figure 5
B
A
C
Figure S5. (A) Comparison of primary cilium incidence in HC (blue), SSc (pink), healthy fibroblasts expressing scrambled shRNA (shSCR, green) and healthy fibroblasts expressing CAV1-targeting shRNA (shCAV1, purple) after serum starvation (n=3). (B) Primary cilia length distribution in HC (blue), SSc (pink), healthy fibroblasts expressing scrambled shRNA (shSCR, green) and healthy fibroblasts expressing CAV1-targeting shRNA (shCAV1, purple) after serum starvation (n=3). (C) Quantification of primary cilium incidence in shSCR fibroblasts (green) and shCAV1 fibroblasts (purple) pretreated with vehicle or MLN8054, followed by treatment with 10 ng/ml TGFβ for 24 h (n=3). All data panels were analysed by ANOVA (ns, non-significant).
